# Leish-ExP: a database of exclusive proteins from *Leishmania* parasite

**DOI:** 10.1101/2020.05.04.076851

**Authors:** Aneesha Das, Nupur Biswas, Saikat Chakrabarti

**Affiliations:** Structural Biology and Bioinformatics Division, CSIR-Indian Institute of Chemical Biology, IICB TRUE Campus, CN-6, Sector 5, Salt Lake, Kolkata, Pin 700091, WB, India

**Keywords:** Leishmania, leishmaniasis, homology, species-specific proteins, genus-specific proteins, drug targeted

## Abstract

Leishmaniasis is a complex disease caused by different species of genus *Leishmania*, which affects millions of people spanning over 88 countries causing extensive mortality and morbidity. Leishmaniasis exists in three clinical forms; cutaneous, mucocutaneous and visceral. Anti-leishmanial therapy is one of the challenging fields due to the presence of wide range of *Leishmania* species and host response against the pathogenic infection. Clinical survey reveals toxicity of drugs and trial compounds causing death among patients and predominant resistance to different class of antimonials among the leishmaniasis patient. Identification of appropriate target protein is a primary and important phase of drug development and *Leishmania* is a suitable system for targeted drug development as it branched out quite early from the higher eukaryotes like human. With the advancement of genomic era and availability of whole genome sequences of *Leishmania* species, relevant information on their respective genes and proteins are now easily extracted. Hence, in the current study, to aid the targeted drug therapy, an exhaustive whole genome scale comparative sequence analysis was performed to identify genes/proteins exclusive to *Leishmania*. Subsequently, further filtration was employed to identify the individual *Leishmania* species specific gene/proteins. Among such proteins, a considerable number was found to have no recognizable structure and functions. Hence, efforts were taken to characterize these *Leishmania* exclusive gene/proteins via providing additional sequence, structure, localization and functional information. All the information about each of these *Leishmania* exclusive gene/proteins from five different *Leishmania* species has been systemically categorized and represented in the form a user-friendly database called **Leish-ExP**, which is freely available at http://www.hpppi.iicb.res.in/Leish-ex.

## Introduction

*Leishmania* is a parasitic pathogen that causes the disease leishmaniasis, more commonly prevalent in the tropical regions. The fact that the disease is endemic and its effect is not manifested in major parts of the world has caused it to be classified as a neglected tropical disease [1]. The disease is spread by the bite of infected sandflies, commonly of the genus *Phlebotomus* in the Indian subcontinent, which serve as vectors for the parasite. Human beings are the primary host of a few of the species of this parasite. Those affected with the disease manifest features as disfiguring scars, lesions on skin besides other disfiguring effects. However, till date there are no proper vaccines available for the leishmanaisis. Chemotherapy, anti-retroviral and anti-carcinoma drugs are mostly used in treating the leishmanial patient. Pentavalent antimonials are the standard first line of treatment but their usage has been severely limited due to emergence of resistance. Chemotherapeutic treatments with amphotericin B, miltefosine and paromomycin are available but their use is limited mostly because of toxicity and/or high cost involvement.

*Leishmania* exists in two different forms within different hosts during its zoonotic life cycle. Sandflies inject infective promastigotes into a susceptible mammal during feeding. Promastigotes are phagocytosed by resident phagocytes, transform into tissue-stage amastigotes, and multiply within these cells through simple division. The parasite continues to infect phagocytic cells either at the site of cutaneous infection or in secondary lymphoid organs, with eventual parasitemia. Sandflies become infected through feeding on a host either with an active skin lesion in CL or with parasitemia in VL. Parasites convert to promastigotes within the sandfly midgut. Promastigotes migrate from the midgut and transform into highly infectious metacyclic promastigotes. In order to survive within the macrophage, which is armed with most potential immune sentinel of human (host) body, parasites are well equipped with outstanding subverting strategies to suppress macrophage immune functions. Multiple parasite derived molecules starting from surface glycolipids like LPG (Lipo Phospho Glycan), GIPL (Glyco Insitol Phospho Lipid) to metal dependent proteases like gp63 and CPC (Cystein peptidases C) have been shown to perform enormous protective activity for amastigotes [2].

Considering the ever-growing resistance and toxicity of the anti-leishmanial drugs, which are primarily non-targeted, it is essential to identify potential new drug targets and subsequently their sets of probable inhibitors. Hence, identification of a pool of genes or proteins that are evolutionarily unique to *Leishmania* and further characterization and categorization of them is due for further target-based drug therapy for leishmaniasis. Since the whole genome sequencing of most of the pathogenic species of *Leishmania sp*. has been completed, a large-scale whole genome comparison allowed us to identify gene/proteins that are exclusive to the *Leishmania* genus. Exclusivity was determined via lack of significant sequence orthology with any other known and sequenced genes/protein. Similarly, *Leishmania* species specific exclusivity was established via lack of significant sequence orthology with any other proteins as well as other *Leishmania* species as well.

We performed exhaustive cross proteome sequence comparison using PSI-BLAST [3] and JACKHMMER [4] programs taking all the sequences from *L. braziliensis, L. donovani, L. infantum, L. major and L. mexicana* and compared each and every one against all other known protein sequences available at the NCBI non redundant (NR) database [5]. Proteins that are exclusively present at the *Leishmania* genus only are termed as *Leishmamia* genus-specific proteins whereas proteins exclusively present in specific *Leishmania* species (e.g., *L. donovani*) are called species-specific proteins. Additionally, we attempt to characterize these *Leishmania* genus and species-specific gene/proteins via providing additional sequence, structure, localization and functional information using *in silico* methods. These *Leishmania* exclusive gene/proteins from five different *Leishmania* species have been systemically categorized and represented in the form a user-friendly database called **Leish-ExP**, which is freely available at http://www.hpppi.iicb.res.in/Leish-ex.

## Methods

### Selection of protein sequences from Leishmania species - L. braziliensis, L. donovani, L. infantum, L. major and L. mexicana

*Leishmania* species were selected based on the availability of their whole genome sequence and their known pathogenicity against human beings. The proteomes of *L. braziliensis, L. donovani, L. infantum, L. major* and *L. mexicana* were obtained from the TriTrypDB database [6]. Protein sequences with special characters such as “*”, “X”, “B” and “Z” were removed from the working dataset. The modified dataset of protein sequences was then subjected to redundancy removal using the standalone version of the CD-HIT software [7]. Further steps were carried out using 8108 proteins from *L. braziliensis*, 7899 proteins from *L. donovani*, 8023 proteins from *L. infantum*, 8044 proteins from *L. major* and 8043 proteins from *L. Mexicana*, respectively. The detailed number of sequences at each step has been shown in **Table 1**.

**Table 1:** Number of *Leishmania* sequences used for comparison.

|  | *Leishmania*<br>*braziliensis* | *Leishmania*<br>*donovani* | *Leishmania*<br>*infantum* | *Leishmania*<br>*major* | *Leishmania*<br>*mexicana* |
| --- | --- | --- | --- | --- | --- |
| **Total number of**<br>**sequences in the**<br>**proteomes** | 8357 | 8032 | 8241 | 8412 | 8452 |
| **Number of sequences**<br>**after removal of**<br>**special characters** | 8197 | 8029 | 8136 | 8330 | 8165 |
| **Number of sequences**<br>**after redundancy**<br>**filter** | 8108 | 7900 | 8023 | 8044 | 8043 |

### Identification of Leishmania genus-specific proteins

The whole proteome of each of the *Leishmania* species was compared against the four others using the homology-scouting software of PSI-BLAST [3] and the results were verified with the help of another software called JACKHMMER [4]. For each of the species, only those protein sequences were considered as genus-specific which had an orthologue in at least one other *Leishmania* species considered in the present study, the maximum being in four other species, but do not have significant sequence similarity with any other protein sequences available in the non-redundant protein (NR) database [5]. Orthology was determined using the E-value threshold of <= 1e^-05^, Query or subject protein sequence length coverage >= 50%, and query and subject sequence identity >= 40%. **Figure 1** shows the flowchart of the protocol used to perform the exhaustive all-to-all proteome level comparison whereas **Tables 2 and 3** show the number orthologous sequences among the five *Leishmania* genomes, retrieved by PSI-BLAST [3] and JACKHMMER [4] programs, respectively.

**Table 2:** Number of orthologous proteins across five *Leishmania* proteomes determined via PSI-BLAST.

|  | Number of<br>orthologous<br>proteins in<br>*L. braziliensis*<br>(8108<br>proteins) | Number of<br>orthologous<br>proteins in<br>*L. donovani*<br>(7900<br>proteins) | Number of<br>orthologous<br>proteins in<br>*L. infantum*<br>(8023<br>proteins) | Number of<br>orthologous<br>proteins in<br>*L. major*<br>(8044<br>proteins) | Number of<br>orthologous<br>proteins in<br>*L. mexicana*<br>(8043<br>proteins) |
| --- | --- | --- | --- | --- | --- |
| *L. braziliensis*<br>(8108 proteins) |  | 7733 | 7883 | 7867 | 7851 |
| *L. donovani*<br>(7900 proteins) | 7533 |  | 7826 | 7735 | 7708 |
| *L. infantum*<br>(8023 proteins) | 7745 | 7918 |  | 7950 | 7935 |
| *L. major*<br>(8044 proteins) | 7757 | 7854 | 7985 |  | 7946 |
| *L. mexicana*<br>(8043 proteins) | 7776 | 7842 | 7979 | 7970 |  |

**Table 3:** Number of orthologous proteins across five *Leishmania* proteomes determined via JACKHMMER.

|  | Number of<br>orthologous<br>proteins in<br>*L. braziliensis*<br>(8108<br>proteins) | Number of<br>orthologous<br>proteins in<br>*L. donovani*<br>(7900<br>proteins) | Number of<br>orthologous<br>proteins in<br>*L. infantum*<br>(8023<br>proteins) | Number of<br>orthologous<br>proteins in<br>*L. major*<br>(8044<br>proteins) | Number of<br>orthologous<br>proteins in<br>*L. mexicana*<br>(8043<br>proteins) |
| --- | --- | --- | --- | --- | --- |
| *L. braziliensis*<br>(8108 proteins) |  | 7790 | 7881 | 7884 | 7874 |
| *L. donovani*<br>(7900 proteins) | 7571 |  | 7827 | 7744 | 7704 |
| *L. infantum*<br>(8023 proteins) | 7786 | 7947 |  | 7940 | 7929 |
| *L. major*<br>(8044 proteins) | 7812 | 7904 | 7986 |  | 7950 |
| *L. mexicana*<br>(8043 number<br>of proteins) | 7831 | 7897 | 7984 | 7964 |  |

**Figure 1:**
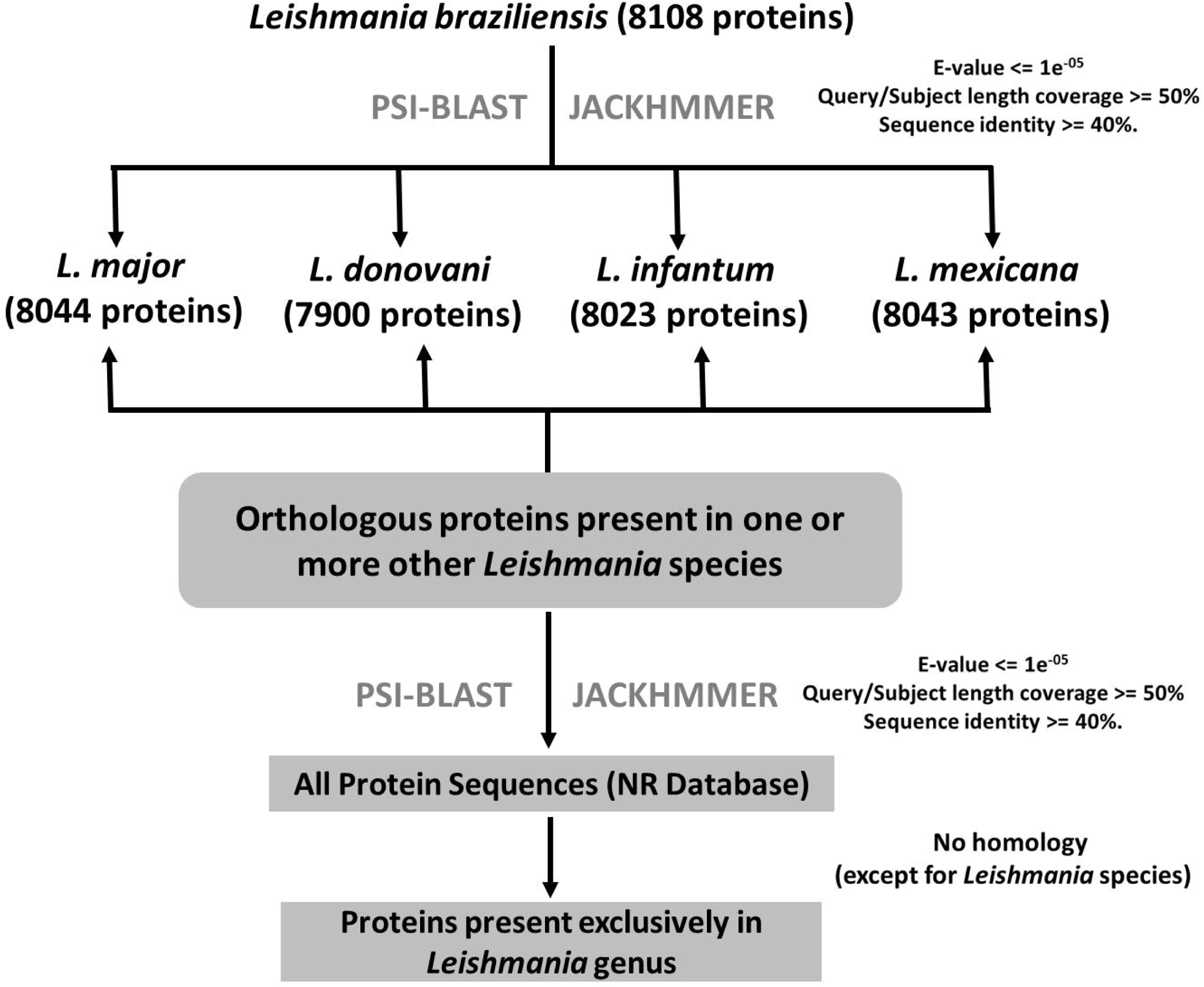
An illustration of the strategy used to identify the genus-specific proteins in each of the *Leishmania* proteome set. This strategy has been shown taking *L. braziliensis* as an example.

### Identification of Leishmania species-specific proteins

Similar to the earlier procedure, whole proteome of each of the *Leishmania* species was first compared against all other four *Leishmania* proteome to identify the orthologous protein sequences between a pair of genuses. Subsequently, the proteins which did not obtain orthologous counterparts with previously described thresholds (E-value: <= 1e^-05^, Query or subject protein sequence length coverage >= 50%, and query and subject sequence identity >= 40%) were subjected to one more round of similarity search against all sequences stored in the non-redundant protein (NR) database. Sequences that fail to obtain any hit passing the selection threshold were considered as specific to particular *Leishmania* species with which the process started. In this way, proteins specific or exclusive to each *Leishmania* species were determined based on sequence orthology. **Figure 2** shows the flowchart of the protocol used to perform the exhaustive all-to-all proteome level comparison whereas **Tables 2 and 3** show the number orthologous sequences among the five Leishmania genomes, retrieved by PSI-BLAST [3] and JACKHMMER [4] programs, respectively.

**Figure 2:**
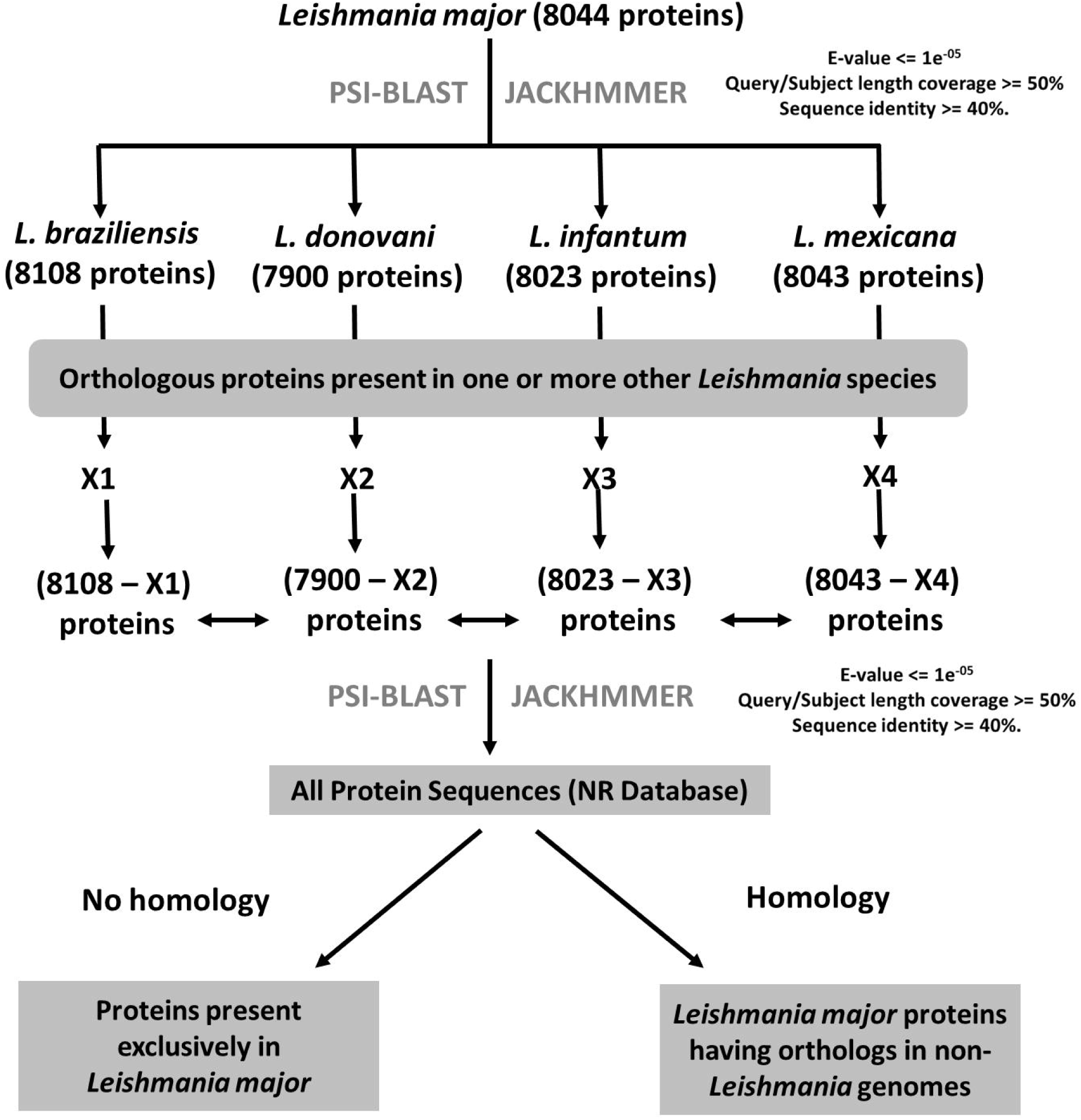
An illustration of the strategy used to identify the species-specific proteins in each of the *Leishmania* proteome set. This strategy has been shown taking *L. major* as an example. X1-X4 are the numbers of proteins from other *Leishmania* species that have orthologous counterpart in *L. major* species.

### Functional annotation of the Leishmania exclusive protein sequences

Functional annotation via homology against protein sequences with known functions stored in the Gene Ontology (GO) database [8] was performed with previously described homology threshold criteria.

Domains refer to those segments within a protein sequence that are involved in performing a biological function as a single unit. Hence, identification of constituent domains within the *Leishmania* exclusive proteins will shed light about their probable functional involvement. The standalone version of the Pfam database [9] was used to scan the Leishmania exclusive protein sequences for the presence of known functional domains.

Sequence motifs are part of a protein sequence that also associates with a particular molecular and/or biological function. The protein sequences, in which known motifs were not previously reported, were subjected to motif search against the PROSITE [10] database using the standalone version of the ScanProsite software [11].

In order to investigate the association of *Leishmania* exclusive sequences with known biological pathways, the Kyoto Encyclopedia of Genes and Genomes (KEGG) [12] were mapped through data mining using in-house developed Perl codes for the purpose.

### Architecture and components of the Leish-ExP database

The Leishmania Exclusive Protein database (**Leish-ExP**) is a repository of protein sequences specific to five *Leishmania* species and genus. The website has been developed in CGI PERL with backend data retrieval from a MySQL database. Following section briefly describes the various parts of the database and its components.

**Leish-ExP** database has two major parts. The first part deals with species specific proteins and associated sequence, structure and functional features whereas the second part deals with genus specific proteins and their functional profiles. Species specific proteins are enriched with the following features

- **Primary sequence analysis:** This tool connects the sequences to obtain information derived from primary sequence using third party tools like, ProtScale [13] and ProtParam [13]. ProtScale is toolbox from ExPASy [14] which provides information regarding bulkiness, molecular weight, polarity, refractivity, number of codons, and probable secondary structural positions. Similarly, another ExPASy tool, ProtParam is also linked to each of the species-specific proteins to provide information regarding its theoretical pI, amino acid composition, atomic composition, extinction coefficient, estimated half-life, instability index, aliphatic index and grand average of hydropathicity (GRAVY).
- **Database retrieval tool:** Links to the sequence databases like TriTrypDB [6], GeneDB [15], and DBGET [16] are also provided for collection additional information regarding the *Leishmania* species-specific proteins.
- **Signal peptide prediction:** Signal peptides are stretches of amino acids located in the protein sequences that direct the protein on their localization in the intracellular milieu. SignalP server [17] implemented in the **Leish-ExP**, predicts the presence and location of signal peptide and cleavage sites utilizing a combination of several artificial neural networks.
- **Sub-cellular localization predictions:** TargetP [18] server utilized in this link to predict subcellular localization of the *Leishmania* species-specific proteins. The localization assignment is predicted based on the presence of any mitochondrial targeting peptide (mTP) or secretory pathway signal peptide (SP). For the sequences predicted to contain an N-terminal pre-sequence a potential cleavage site can also be predicted.
- **Prediction of MHC binding peptide:** Sequence based analysis has also been done for species-specific proteins to predict whether they contain any peptide sequence that can bind to major histocompatibility complex (MHC) molecules. For this purpose, results from the Rankpep [19] program have been represented as potential binding peptides to Class I and Class II MHC molecules.
- **Prediction of phosphorylation sites:** Identification of probable phosphorylation sites in proteins is an important aspect as it provides critical insight towards the function of the protein. NetPhos 2.0 server [20] was used which uses neural network-based predictions of serine, threonine and tyrosine phosphorylation sites.
- **Prediction of secondary structure:** The knowledge about the structures of proteins is important to determine their functions. In absence of the tertiary structure, information regarding the secondary structures of a protein is critical for understanding its biophysical and biochemical properties. **Leish-ExP** provides residues wise secondary structures (α helix, β strand, and coil) predicted by PSI-PRED [21] for each *Leishmania* species-specific protein.
- **Computation of physico-chemical properties:** Physico-chemical properties of a protein are critical factors in determining its unique shape, size, stability, biochemical and biological functions. Similarly, it is also known that the presence or absence certain functional groups such as AMDO (Gln and Asn), AMNP (Lys), CBXL (Asp and Glu), GNDO (Arg), HDXL (Ser, and Thr), IMZL (His), NONP (Ala, Gly, Ile, Leu, Val and Pro), PHEN (Phe, Trp and Tyr), SULF (Met), THIO (Cys) at the protein surface or code can influence the function of the protein greatly. Hence, fractions of these functional groups are also calculated for each of the species-specific protein in **Leish-ExP** database and are represented in graphical format.

The second part of the **Leish-ExP** database lists the genus-specific proteins and their associated sequence, structural and functional features. Additionally, this part also categorizes genus-specific proteins based on their functional classification. All the above mentioned toolbox such as, primary sequence analysis, database retrieval tool, signal peptide prediction, Sub-cellular localization predictions, prediction of MHC binding peptide, prediction of phosphorylation sites, prediction of secondary structure, computation of physico-chemical properties and their analytical results are also provided for *Leishmania* genus-specific proteins as well. Functional annotation of the genus-specific proteins was performed using the Gene Ontology (GO) database with previously described homology threshold criteria. However, only molecular function categories broadly divided into enzyme and non-enzyme class was presented in the **Leish-ExP** database.

In addition to the above components, the **Leish-ExP** database has two other useful options where a user can search the species-specific and genus-specific proteins via keyword based *‘Search’* option. The option entitled *‘Server’* allows the user to run BLASTp [3] similarity search against the *Leishmania* proteomes as well as against the *Leishmania* genus-specific proteins.

## Results

### Leishmania genus-specific and species-specific proteins

5855 genus-specific and 183 species-specific proteins were identified using the above-mentioned protocol and criteria (see Methods). Interestingly, a significant number of proteins were found to be restricted within the *Leishmania* genus (∼23% of the combined *Leishmania* proteomes, **Table 4**) whereas a much lower number were found to be individual species-specific (**Table 4**).

**Table 4:**
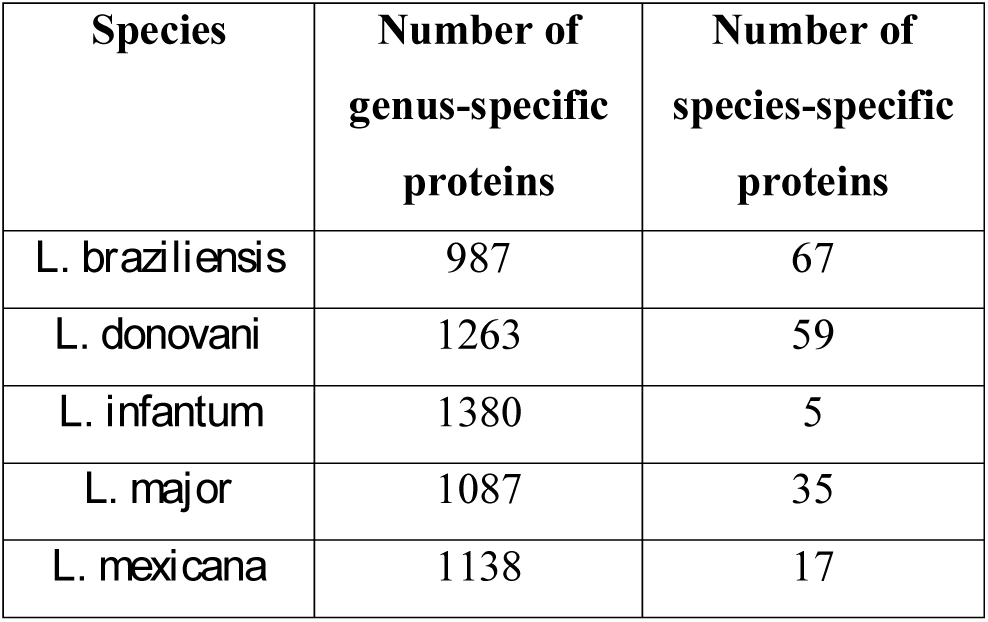
Number of *Leishmania* exclusive proteins.

### Functional assignment of the Leishmania exclusive protein sequences

Functional assignment was done to *Leishmania* genus-specific proteins out of which 169 were assigned to various enzyme class and 190 to non-enzyme class. **Figure 3** shows the distribution of various enzymes and non-enzymatic functional categories within these proteins.

**Figure 3:**
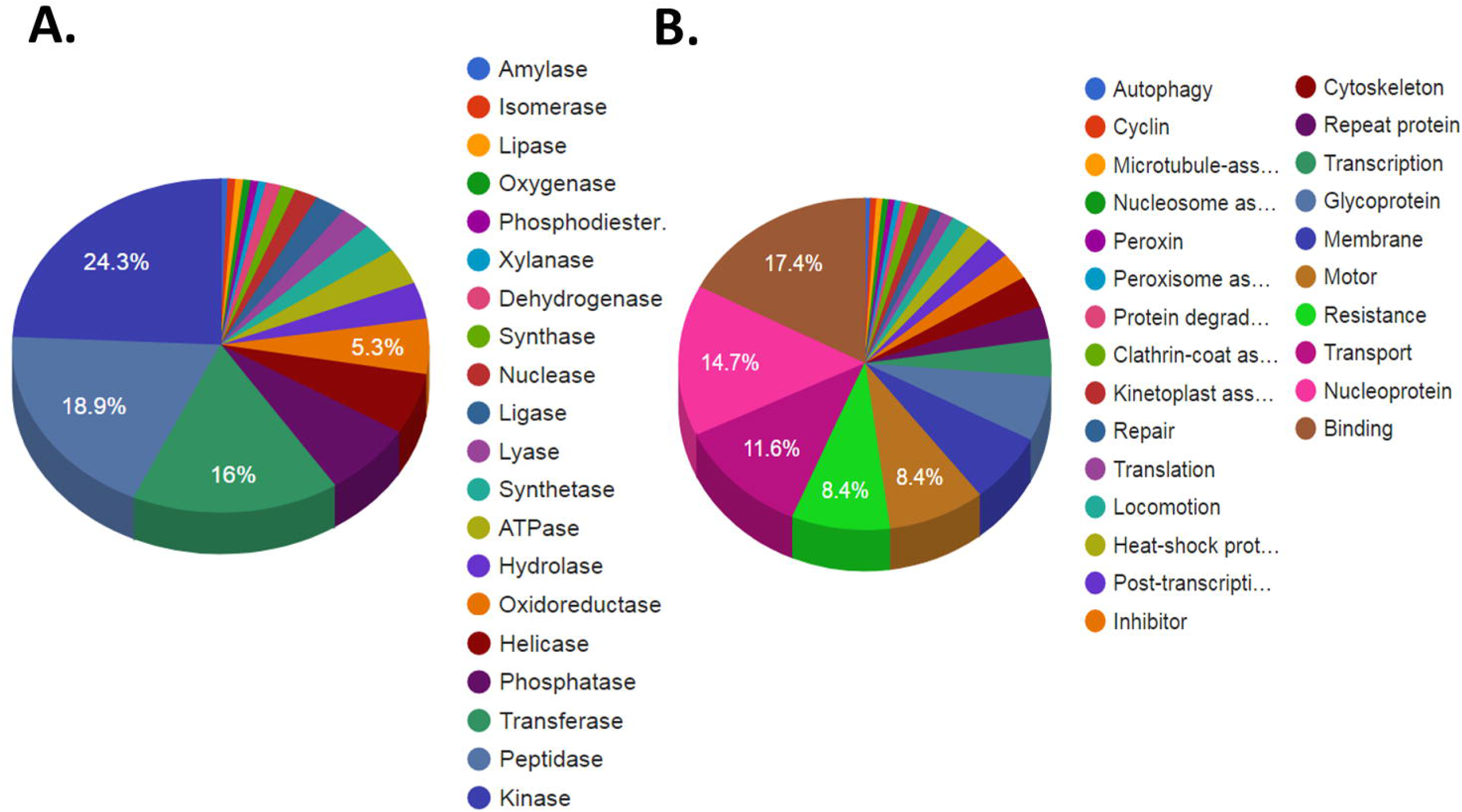
Functional distribution of the *Leishmania* genus-specific proteins that were annotated using sequence orthology based approach against GO database for enzyme (A) and non-enzyme (B) categories.

Domains are certain regions within the protein that are capable of folding in an independent manner and also can perform individual functional role. In absence of three-dimensional structure for the genus and species specific *Leishmania* proteins, detection of domains provided important leads to the detection of function of these proteins. The sequences were scanned against the Pfam [9] domains database using the hmmscan [4] algorithm. Those sequences that had a homolog with an e-value of less than or equal to 1e-05, subject coverage of greater than or equal to 50% and an identity of greater than or equal to 40% were considered for domain assignment. Quantitative analysis suggests that only a limited number of domains could be assigned to the genus-specific proteins. However, most frequently mapped domain within the *Leishmania* genus-specific proteins are MORN (Membrane Occupation and Recognition) domain, Zinc finger domain, Leucine-rich repeat domain, RING finger domain, Proline-Glycine-Proline repeat domain, ATPase domain, WD40 repeat domain, and Kinesin motor domain are few of them.

In the present study, the genus-specific protein sequences as well as the species-specific sequences were scanned against the Prosite database [10] using the ScanProsite [11] algorithm to find significant match with known motifs. Numbers of *Leishmania* genus-protein sequences annotated by motif mapping are 57, 111, 143, 73 and 92 for *L. braziliensis, L. donovani, L. infantum, L. major and L. mexicana*, respectively. Most frequently observed motifs are protein kinase motif, calmodulin-binding IQ motif, TPR-repeat motif, Leucine-rich repeat, and EF-hand calcium-binding motif.

Pathway mapping against the KEGG database could map 29 genus-specific proteins with known pathways.

### Comparison of physico-chemical properties

The functions that are performed by proteins are attributed largely to their physico-chemical properties. For instance, whether a protein will be acidic or basic is dictated by the kind of amino acids that are predominant in the sequence. The Amino-Acid-Index database [22] is a repository that contains a collection of physico-chemical property indices calculated for each amino acid for all 544 physico-chemical properties listed in the AAI database. In this study, the indices were recruited to decipher the score for a particular physico-chemical property for each protein sequence. This was done by multiplying the index value representing an amino acid for a particular physico-chemical property by the number of that particular amino acid in the protein sequence. The score for the entire protein sequence was established by summing up the products of the index and the number of amino acids computed in the previous step, which was then divided by the number of amino acids in the protein sequence to obtain the average score for that particular protein. This score was used to compare any difference between the scores of the genus-specific proteins and other leishmanial proteins (excluding the species-specific proteins), as well as between the species-specific proteins and other proteins (excluding the genus-specific proteins). Before proceeding with the study, the *Leishmania* exclusive proteins were categorized into four groups (short, medium, long and very long) according to the lengths of the sequences. The above steps were taken to eliminate any discrepancy arising out of comparison done among protein sequences which had large differences in lengths. The R program (version 3.1.3) [23] was used to compute the difference between sets of proteins in the same category by means of Welch’s t-test [24] using the following formula:

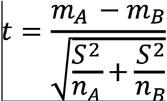

where, *m*_*A*_= mean value for the parameter from the genus-specific/species-specific set of proteins,

*m*_*B*_ = mean value for the parameter from other set of proteins,

*S* = standard deviation,

*n*_*A*_ = degrees of freedom for the genus-specific/species-specific set of proteins,

*n*_*B*_ = degrees of freedom for the parameter from other set of proteins.

The physico-chemical properties between the genus-specific protein sequences and other proteins (excluding the species-specific proteins) showing significant differences are tabulated in **Table 5** whereas **Table 6** provides the significantly different physico-chemical properties between the species-specific protein sequences and other proteins (excluding the genus-specific proteins).

**Table 5:** Significantly different physico-chemical properties between *Leishmania* genus-specific and other remaining proteins.

| **Sequence length category** | **Significantly different physicochemical properties** | **Property status in genus-specific proteins** |
| --- | --- | --- |
| Short | C-cap residue (GLY) frequency at helix C terminal | LESSER |
| Medium | Hydrophobicity | LESSER |
| Long | Hydrophobicity | LESSER |
| Very long | C-cap residue (GLY) frequency at helix C terminal | LESSER |

**Table 6:**
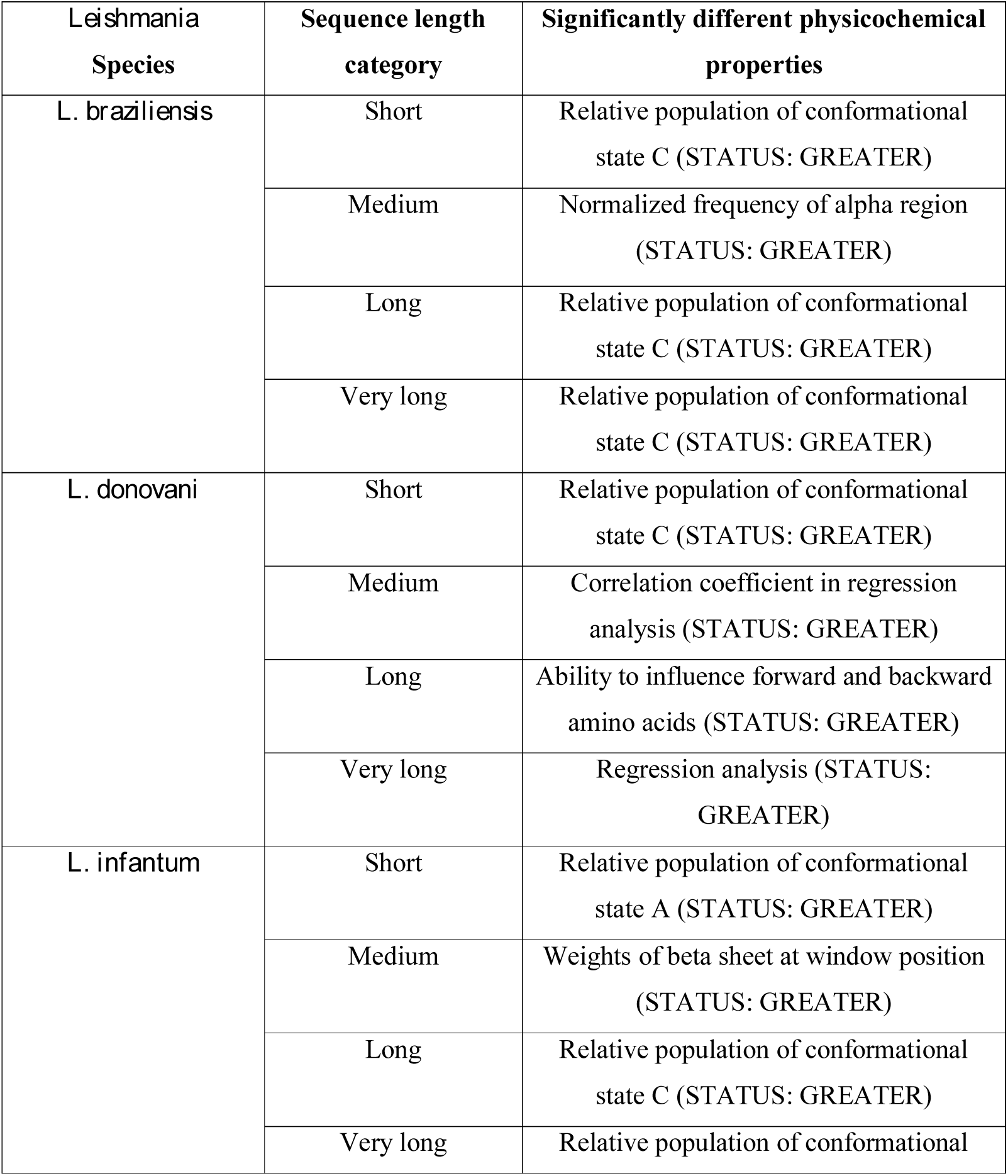
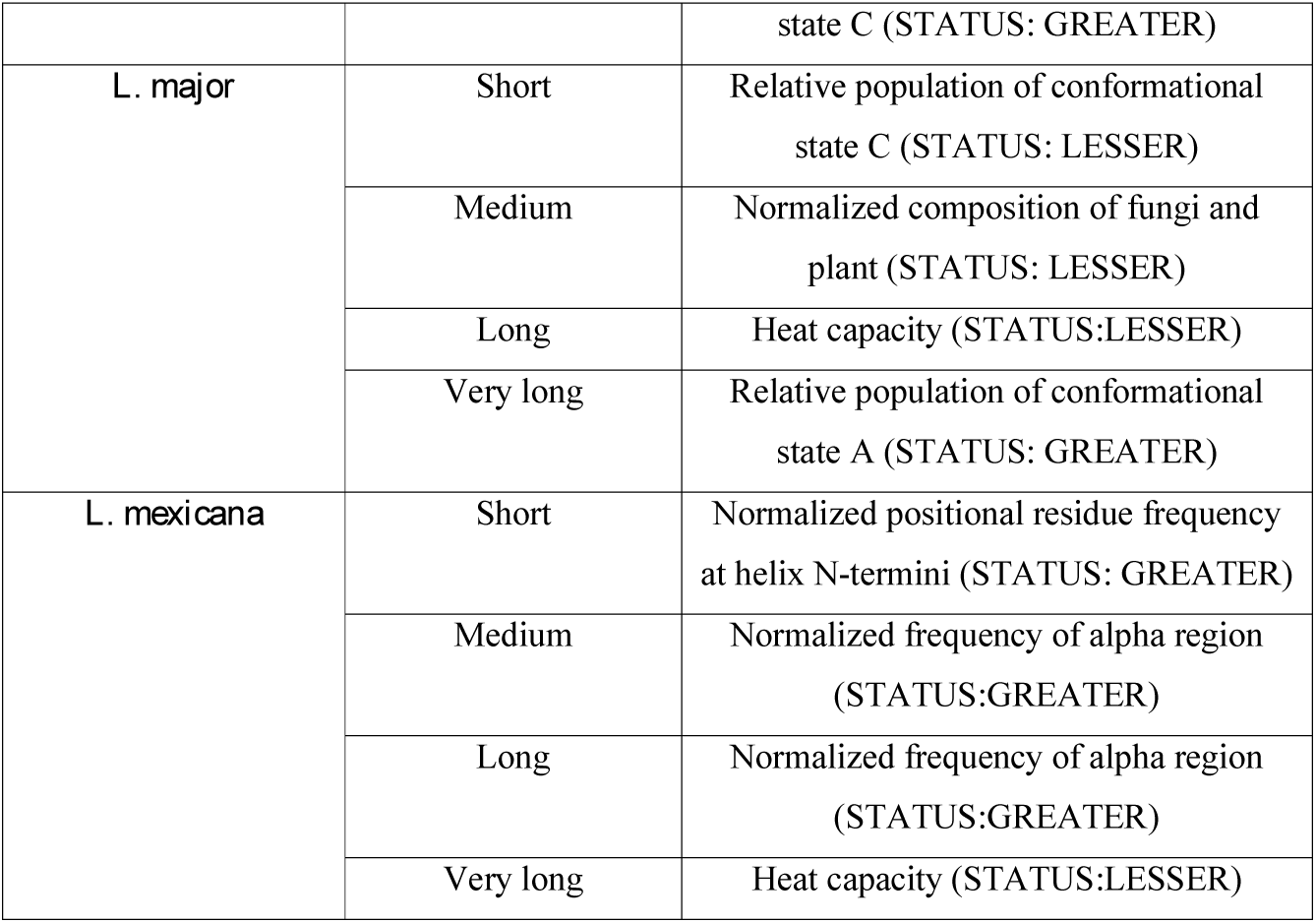
Significantly different physico-chemical properties between *Leishmania* species-specific and other remaining proteins.

### Leish-ExP database

The **Leish-ExP** database lists the identified species and genus specific proteins and also provides additional information regarding the primary sequence and its attributes, physico-chemical properties of the constituent amino acids, sub-cellular localization, secondary structure, and probable functional association. Detailed description of the individual components has been discussed in the Methods section. **Figure 4** provides a snapshot of the analyses done by various tool box of the **Leish-ExP** database for the species-specific proteins. These are displayed by clicking on the tab with the name of the *Leishmania* species on the home page.

**Figure 4:**
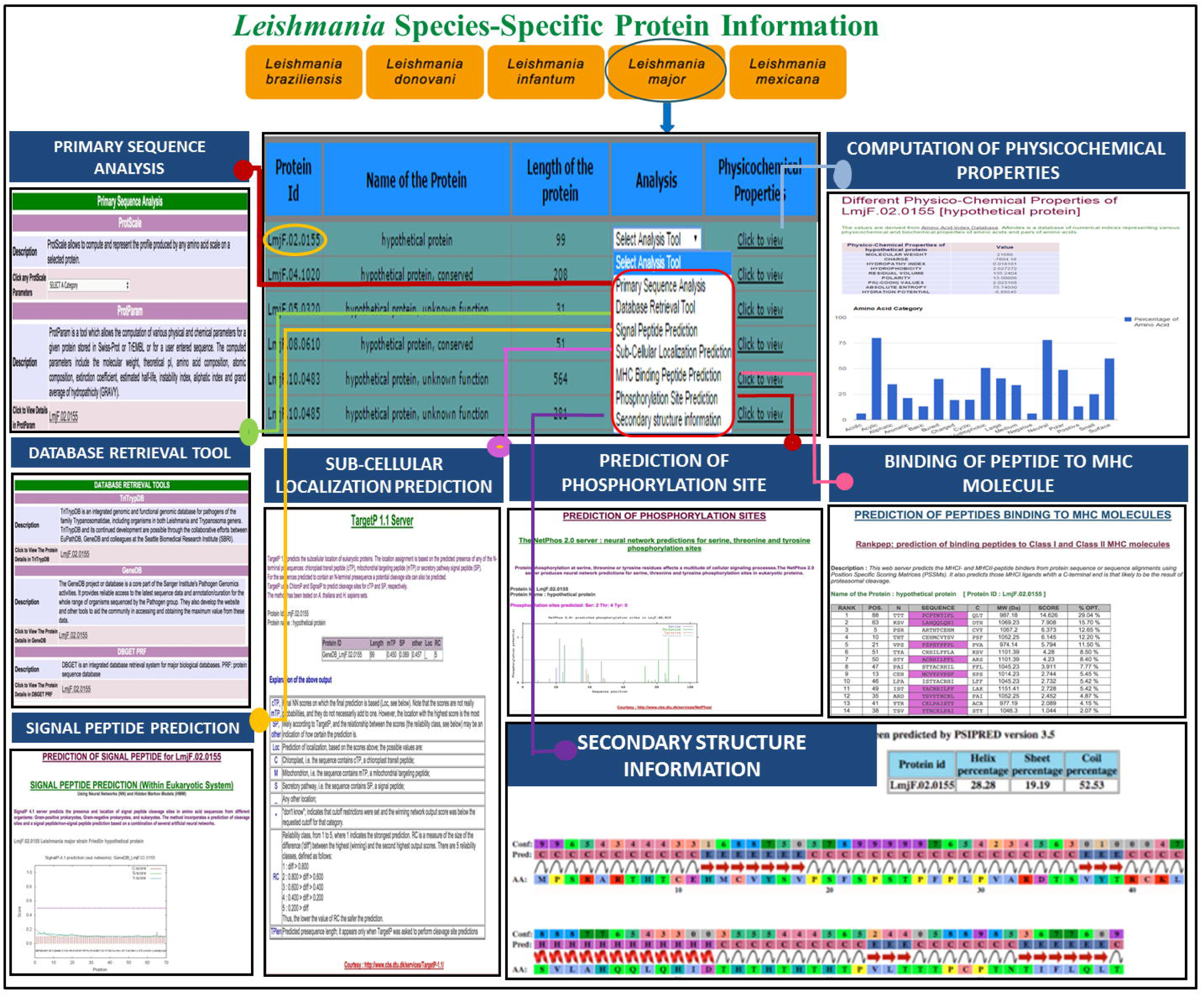
A combined snapshot of the various features of the *Leishmania* species-specific proteins in **Leish-ExP** database.

## Discussion

Leishmaniasis is a vector-mediated disease prevalent worldwide by various types of *Leishmania* species with broad spectrum of clinical manifestations, ranging from self-healing cutaneous lesions to visceral leishmaniasis (VL). Some of the Leishmaniasis manifestation has not only severe impact on global health and economy but also leaves a profound social stigma [25]. In addition, the fatal form of leishmaniasis, VL still remains to be one of the top parasitic diseases with outbreak and mortality potential [26]. Toxicity of the existing drugs and continuous emergence drug resistance among the leishmaniasis patients is a major threat to the existing disease control and treatment regime and perhaps can be associated with development of secondary ailments such as post-kala-azar dermal leishmaniasis (PKDL), which is usually a sequel of visceral leishmaniasis that appears as macular, papular or nodular rash usually on face, upper arms, trunks and other parts of the body.

*Leishmania* specific databases are not new. There are generic databases like GeneDB [15] and TriTrypDB [6], which provides genomic and genome sequencing derive information. There are a few specific databases as well, such as The LeishCyc database [27] detailing the metabolic pathways and metabolite relationship of the *Leishmania major* genes. Similarly, LeishDB is a database for storing data related to coding genes and non-coding RNAs from *L. braziliensis* [28]. LmSmdB [29] is an integrated database for metabolic and gene regulatory network in *Leishmania major* while Leish-ESP [30] database enlist *Leishmania* gene expression and transcriptome information. However, to the best of our knowledge, there is no database available which identify, categorize, and characterize genes/proteins exclusive to *Leishmania* genus and individual species. Hence, **Leish-Exp** database provides unique and useful information which could be utilized further for therapeutic target design in leishmaniasis.

Targeted drug therapy provides a far better control and precision over non-targeted anti-microbial drugs, especially for the ones that are known to develop resistance via modification of their genome sequences. Hence, identification of novel and druggable target proteins are part of a successful targeted drug designing scheme. In order to address this issue, we collected a pool of genes or proteins that are evolutionarily unique to *Leishmania* genus and five major disease-causing species. We also attempted to characterize these *Leishmania* exclusive proteins and categorize them in a way so that the information becomes helpful for further target-based drug design. Here, we performed a large-scale whole genome comparison to identify gene/proteins that are exclusive to the *Leishmania* genus and species via implementation of sequence homology based search. Exclusivity was determined via lack of significant sequence orthology with any other known and sequenced genes/protein. These *Leishmania* exclusive gene/proteins from five different *Leishmania* species have been systemically categorized and represented in the form a user-friendly database called **Leish-ExP** freely available at http://www.hpppi.iicb.res.in/Leish-ex.

## Authors’ contributions

SC conceptualized the project. AD and NB designed the web server. AD performed the analysis. SC analyzed the data and drafted the manuscript.

## Acknowledgement

The authors acknowledge CSIR-Indian Institute of Chemical Biology for infrastructural support. SC acknowledges CSIR Network Project (BSC-0114) and the Systems Medicine Cluster (SyMeC) grant (GAP357), Department of Biotechnology (DBT) for funding. NB acknowledges the Systems Medicine Cluster (SyMeC) grant (GAP357), Department of Biotechnology (DBT) for fellowship. AD acknowledges Council of Scientific and Industrial Research (CSIR) for fellowship.

